# Motor unit discharge properties are modestly influenced by menstrual cycle-related fluctuations in sex hormone concentrations

**DOI:** 10.64898/2026.01.16.699975

**Authors:** Sophia T. Jenz, Pádraig Spillane, Mollie O’Hanlon, Elisa Nédélec, The MUSH Collaboration, C.J. Heckman, Mathew Piasecki, Paul Ansdell, Jessica Piasecki, Gregory E.P. Pearcey

**Author notes:** Address for correspondence: Gregory Pearcey, School of Human Kinetics and Recreation Memorial University of Newfoundland, St. Johns, NL, Canada, A1C 5S7. Authors with equal contributions (Sophia T. Jenz, Pádraig Spillane). In addition to the authors named above, the MUSH Collaboration includes the following contributors (listed in alphabetical order with their specific contributions as per the National Information Standards Organization [NISO] Contributor Roles Taxonomy). James A. Beauchamp^6^: Formal analysis; Software; Writing – review & editing Alex Benedetto^1,7^: Investigation; Writing – review & editing Melissa E. Fajardo^1^: Investigation; Writing – review & editing Colin K. Franz^7,8^: Project administration; Resources; Writing – review & editing Stuart Goodall^2^: Supervision; Methodology; Writing – review & editing Kirsty M. Hicks^2^: Supervision; Methodology; Writing – review & editing Tea Lulic-Kuryllo^9,10^: Methodology; Writing – review & editing Francesco Negro^11^: Formal analysis; Software; Writing – review & editing Elisa Pastorio^2^: Investigation; Writing – review & editing Anna Wigbers ^12^: Investigation; Writing – review & editing.

## Abstract

Concentrations of estradiol (E2) and progesterone (P4), the main female sex hormones, exhibit large fluctuations across the menstrual cycle. Due to their receptors throughout the central nervous system, both hormones have the potential to influence motor function by influencing ionotropic and metabotropic inputs to motor pools, which can be estimated through the neural codes extracted from motor unit discharge patterns. To address key methodological limitations in prior menstrual cycle research on motor output, we established the Motor Units and Sex Hormones (MUSH) collaboration. The objective of this multi-site investigation was to determine whether endogenous fluctuations in estradiol and progesterone influence human motor unit activity. We hypothesized that motor unit discharge rates and persistent inward current (PIC)–related contributions to discharge would be greatest during the late follicular phase, when estradiol concentrations were highest. Fifty females completed a comprehensive protocol involving menstrual cycle and ovulation tracking, serum hormone measurement, and high-density surface electromyographic recordings during isometric contractions to quantify motor unit activity in the early follicular, late follicular, and mid luteal phases. After exclusion of 10 females with either atypical hormone concentration profiles or insufficient motor unit yield, 40 remained in the final analysis. There were significant changes in several motor unit discharge variables between menstrual cycle phases and significant associations with hormone concentrations. Increased estradiol was associated with higher peak discharge rates and ascending discharge rate nonlinearity, while increased progesterone was associated with higher peak discharge rates, more discharge rate hysteresis and ascending discharge rate nonlinearity. Despite reaching statistical significance, the magnitudes of these effects (i.e., effect sizes) were small. Overall, these findings indicate that fluctuations in sex hormones influence motor unit behavior, but the effects are subtle, highlighting the need for well-powered and methodologically rigorous menstrual cycle research.

**Graphical Abstract:** 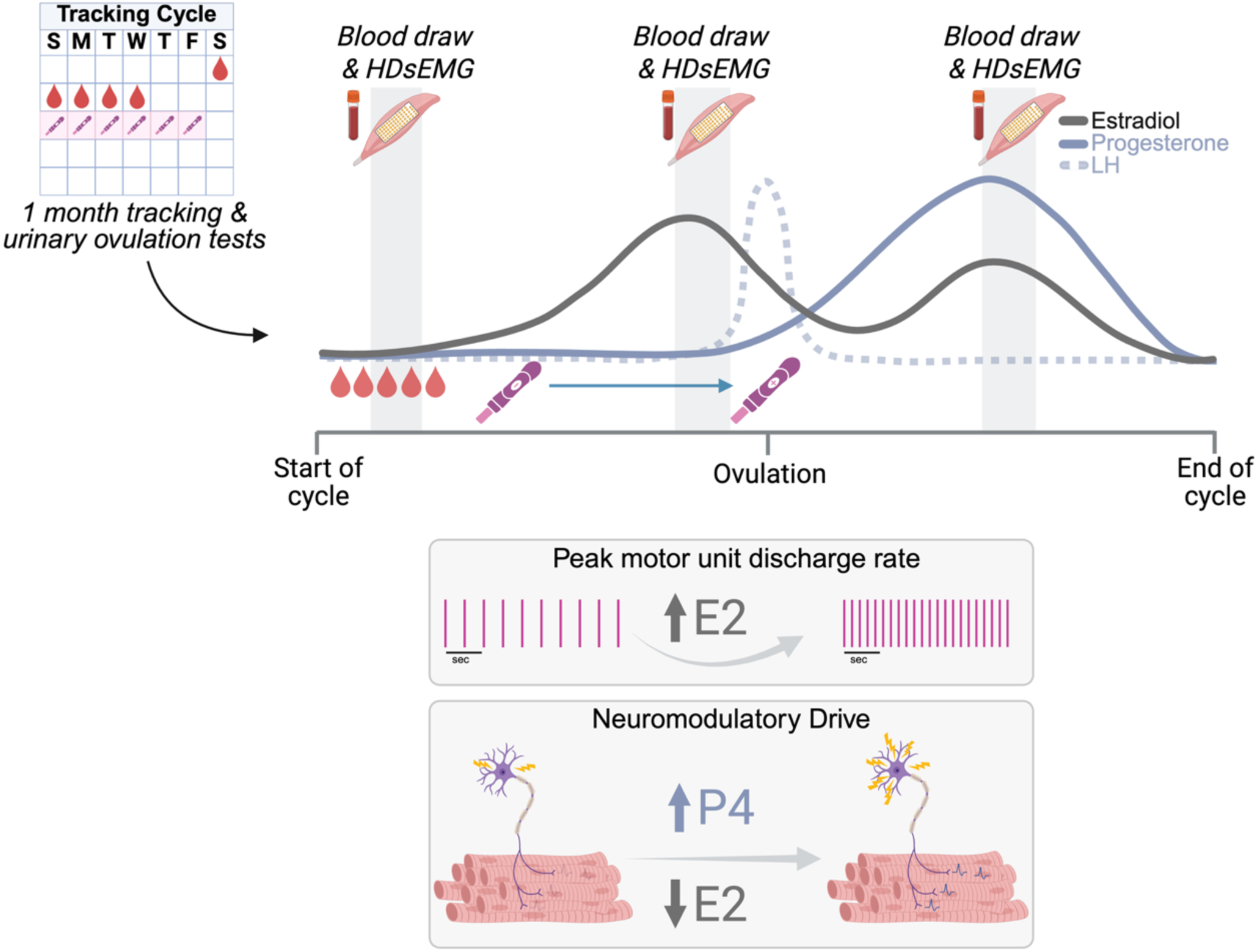

**KEY POINTS:**

1. There are small but detectable differences in motor unit discharge rates between menstrual cycle phases, which are predicted by within-participant fluctuations in estradiol and progesterone.
2. Discharge rate patterns that provide estimates of neuromodulatory and inhibitory input suggest that estradiol and progesterone can influence spinal cord circuitry differently than has previously been documented in the brain, highlighting an understudied aspect of female neurophysiology.
3. Variability in menstrual cycles and associate hormones makes large-scale, rigorous studies especially valuable in female neuromuscular research.

## INTRODUCTION

Recent evidence indicates that menstrual cycle-related fluctuations in sex hormones might influence female physiology, disease, and nervous system function (Roeder & Leira, 2021; Rugvedh *et al*., 2023; Mairajuddin *et al*., 2025). Since the nervous system underlies the control of movement, sensation, and pain, it is critical to identify whether its function differs across the menstrual cycle. However, investigating whether the neural control of movement is altered across the menstrual cycle remains challenging due to the high inter- and intra-individual variability in cycle characteristics (Fehring *et al*., 2006), the cost and complexity of hormonal measurements, and the underrepresentation of females in research (Costello *et al*., 2014; Lulic-Kuryllo & Inglis, 2022; Ranadive & Hagberg, 2025). Neurophysiology and exercise science investigations on the menstrual cycle are often under powered and use inconsistent methods for phase identification, limiting the validity, reproducibility, and interpretability of findings (Elliott-Sale *et al*., 2021; Schlie *et al*., 2025). To address the limitations of small samples and hormonal verification, we established a multi-site collaboration to study relationships between motor units and sex hormones (MUSH). This international effort aims to generate a robust dataset that advances the physiological understanding of the female motor system.

Motor units are comprised of spinal motoneurons and their innervated muscle fibers, which form the final common output of the motor system (Heckman & Enoka, 2012). Due to the one-to-one ratio of action potentials with their innervated muscle fibers, motoneurons are the only cell that can be routinely and non-invasively recorded from the human central nervous system. High-density surface electromyography (HDsEMG) is a robust tool that allows for the identification of motor unit spike instants, which are responsible for movement control that is achieved with two main strategies: motor unit recruitment and rate coding (Adrian & Bronk, 1929). Both strategies work in unison and rely on a dynamic balance of excitatory, inhibitory, and neuromodulatory inputs (Heckman *et al*., 2009). With recent advances in HDsEMG, a clearer understanding of how these inputs change across tasks and effort levels is developing (Škarabot *et al*., 2025); specifically, how the interactions between neuromodulatory drive and patterns of inhibition sculpt motor output (Beauchamp *et al*., 2023; Chardon *et al*., 2024). Such data provides rich information about the motor system and allows for simultaneous examination of cellular mechanisms that may underlie fluctuations in female motor output.

Although females remain understudied in motor unit studies, sex-related differences have become evident in recent years (Lulic-Kuryllo & Inglis, 2022). Yet it remains unclear whether differences arise from intrinsic motoneuron properties, descending inputs, muscle contractile characteristics, or a combination of factors (Jenz & Pearcey, 2022). Unlike males, females experience substantial fluctuations in sex hormones across the menstrual cycle (Sherman & Korenman, 1975). Two of these sex hormones, estradiol (E2) and progesterone (P4), cross the blood-brain barrier and can have broad effects on the central nervous system (Bernal & Paolieri, 2022). In humans, this has been shown to impact cortical and corticospinal excitability, with estradiol having neuro-excitatory effects, and progesterone exerting inhibitory effects (Smith *et al*., 2002; Ansdell *et al*., 2019; Spillane *et al*., 2026). Despite some evidence to support sex hormone effects on neural excitability, most work at the motor unit level is often underpowered and lacks consistency when defining menstrual cycle phases, resulting in data that is not reproducible and difficult to interpret.

There is a strong rationale to explore how sex hormones affect motoneurons via the monoaminergic systems. Serotonin and norepinephrine both facilitate persistent inward currents (PICs; Lee & Heckman, 1998, 2000), and both estradiol and progesterone have been shown to enhance serotonergic signaling in reduced preparations (Sumner & Fink, 1993; Gundlah *et al*., 2000; Bethea *et al*., 2002; Centeno *et al*., 2007; Rivera & Bethea, 2012), which has the capacity to increase motoneuron excitability. When considered alongside evidence of plateau potentials during estrus in anesthetized cats (Kirkwood *et al*., 2002; Ford & Kirkwood, 2006), these data provide a plausible mechanism for the recently observed sex differences in estimates of PIC magnitude in humans. Our groups have recently shown that females generally have higher motor unit discharge rates than males (Piasecki *et al*., 2021; Guo *et al*., 2022; Jenz *et al*., 2023), which are accompanied by larger estimates of PICs (Jenz *et al*., 2023; Lecce *et al*., 2024; Yacyshyn *et al*., 2025). However, the hormonal contributions to these differences are not widely understood.

To determine whether endogenous estradiol and progesterone concentrations influence motor unit activity, we used HDsEMG to noninvasively quantify motor unit discharge rates and estimate PIC-related contributions to these discharge patterns in the tibialis anterior (TA) across three hormonally distinct phases of the menstrual cycle. Based on estradiol’s known role in upregulating serotonergic signaling in the central nervous system, we hypothesized that discharge rates and estimates of PIC contributions to motor unit output would be greatest during the late follicular phase, when estradiol peaks. The findings from the first MUSH collaboration study are presented in this manuscript.

## METHODS

### Participants and menstrual cycle information

#### Participants

This study focuses on the effects of the dominant female sex hormones on motor unit discharge behavior during submaximal isometric contractions. The MUSH collaboration defines *female* as individuals assigned female at birth (Institute of Medicine (US) Committee on Understanding the Biology of Sex and Gender Differences, 2001). It is important to note that while we investigate physiological differences in biological sex and the effects of sex hormones, we acknowledge that gender identity and environment also influence human physiology. Fifty females volunteered to take part in this multi-site research study. Participants reported no history of neurological injury or disease, no musculoskeletal injury in their lower right limb, regular menstrual cycles (≥ 21 and ≤ 35 days), and no hormonal contraception use or pregnancy within the last 12 months. All participants provided written consent to the protocol which was approved by each institutional review board at the respective sites. General methodology was kept consistent across all three sites but details unique to each university are described in Supplemental Table 1.

### Cycle tracking and scheduling

One cycle before scheduling sessions, participants reported the days of menses and tracked predicted ovulation using urine luteinizing hormone (LH) test strips (**Figure 1A**). Sessions were scheduled in the subsequent cycle(s) as accurately as possible to capture three key phases of the menstrual cycle: early follicular (EF; during menses with a test window of cycle day 2 to 4), late follicular (LF; 24 to 48 hours prior to LH surge), and mid luteal (ML; 7 to 9 days from a positive urinary LH test; **Figure 1B**). The session order was randomized, with some participants completing session one in the early follicular, late follicular, or mid luteal phase. Each session was scheduled at the same time of day. During the experimental cycles, participants also tracked and reported the day of LH surge. If at any point a participant reported no ovulation for two consecutive cycles, they were excluded from the remainder of the study.

**Figure 1.**
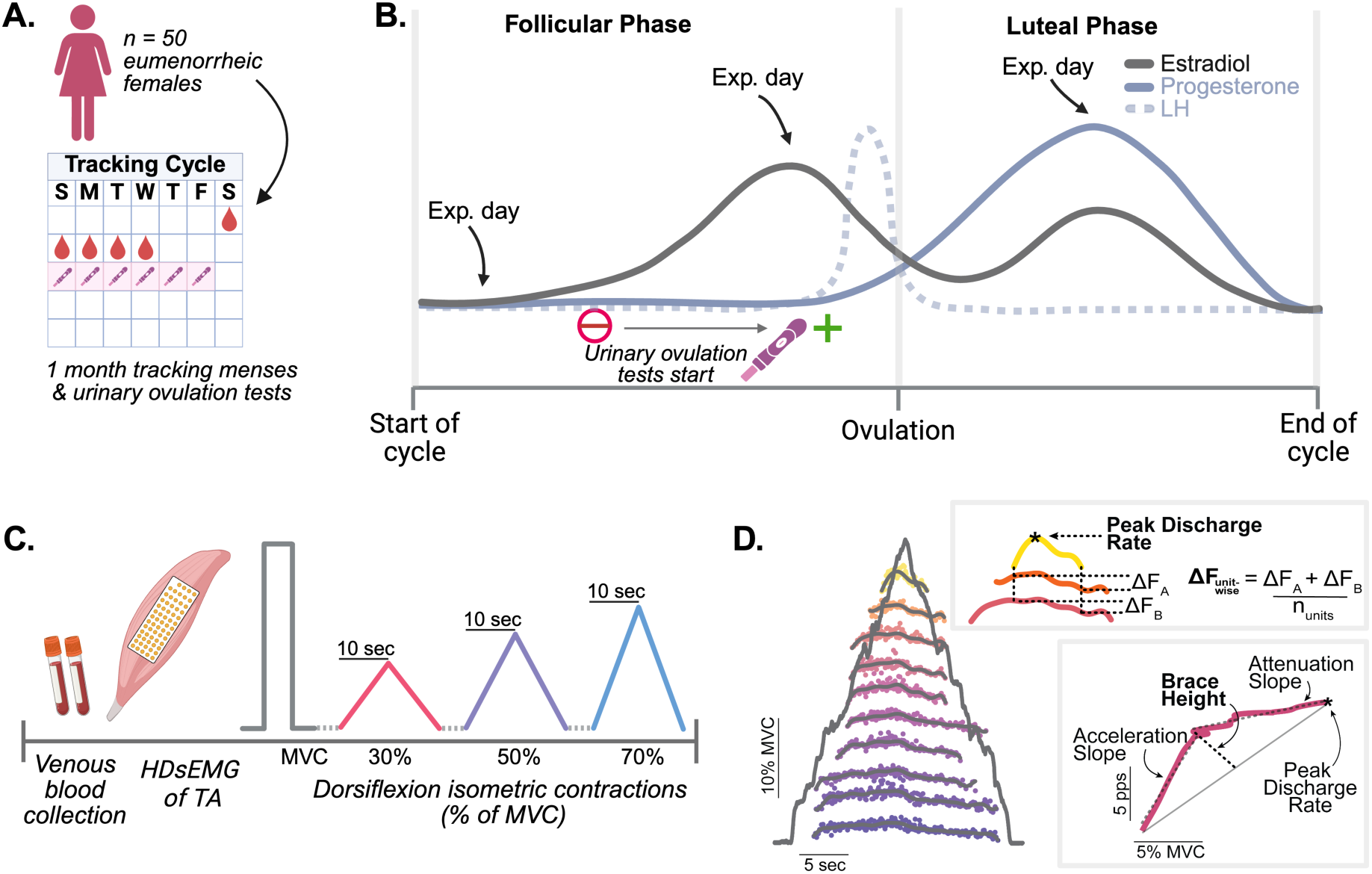
Methodology across all three sites. (A) One month of menstrual cycle and ovulation tracking prior to the experimental sessions. (B) Illustration of the theoretical hormonal fluctuations and the scheduling of early follicular, late follicular, and mid luteal sessions across a cycle. (C) Overview of the in-session HDsEMG recording protocol. (D) Motor unit post-processing workflow, including smoothing of discharge rates and calculation of discharge rate hysteresis (ΔF) and ascending discharge rate nonlinearity (brace height).

### Hormone analysis and exclusion

At the beginning of each session, participants received a blood draw from a trained phlebotomist to measure serum estradiol and progesterone concentrations. Hormone concentrations were used to retrospectively confirm cycle status and for subsequent analysis (**Figure 1C**). Participants were excluded if progesterone concentration was lower than 12 nmol/L in mid luteal phase (NHS North Bristol Trust, 2025) or if there was no positive ovulation test during the testing cycle.

#### In Session Protocol

Participants were seated with their ankle secured in a dynamometer at 90-100°, with a monitor in front of them to provide visual torque feedback. Torque was sampled at 2,048 Hz and smoothed offline with a fifth-order Butterworth filter. Participants began each session by performing isometric dorsiflexion maximum voluntary contractions (MVC), where they received verbal encouragement to ensure maximal effort (**Figure 1C)**. One minute of rest was given between each trial, and if the final trial had the highest MVC value, additional trials were performed. The MVC trials were used to normalize the rest of the experimental trials. Participants performed triangle contractions with a 10 second up and down phase to 30, 50, and 70% of their MVC in a randomized order (**Figure 1C)**. A minimum of two ramps per intensity were performed, but trials were discarded and repeated if the torque trace showed substantial deviations from the target ramp.

### High-density surface electromyography

A high-density surface electromyography (HDsEMG) electrode grid (64 electrodes, 13 x 5 configuration, 8 mm inter-electrode distance, GR08MM1305, OT Bioelettronica, Turin, Italy) was placed over the TA. HDsEMG signals were recorded in monopolar mode, sampled at 2048 Hz, amplified 150 times, and band-pass filtered between 10-500 Hz using a signal amplifier (OT Bioelettronica, Turin, Italy). Further site specific HDsEMG methods can be found in Supplemental Table A1.

### Data Analysis

#### Motor unit decomposition

EMG signals with substantial artifacts or noise were removed, and individual motor unit spike trains were decomposed using a convoluted blind-source separation algorithm, after which motor unit spike trains were manually edited to correct minor errors made by the decomposition (see Supplemental Table 1 for site specific tools). Using custom MATLAB scripts, motor unit instantaneous discharge rates (IDRs) were calculated using the inverse of the interspike interval, and spike trains were smoothed using support vector regression (Beauchamp *et al*., 2022). From the smoothed spike trains, initial, peak, and final discharge rates were calculated. Motor unit recruitment and derecruitment thresholds were quantified as the percentage of MVC at the time of the first and final spike.

### Motor unit discharge rate pattern analysis

The contributions of PICs to discharge were estimated by quantifying both discharge rate hysteresis and ascending nonlinearity. Discharge rate hysteresis was calculated using the unit-wise paired motor unit analysis (i.e., ΔFrequency, or ΔF), as previously validated by our lab and others (Gorassini *et al*., 2002; Powers & Heckman, 2015) and described in prior work (Jenz *et al*., 2023). A mean unit-wise ΔF was calculated for each test unit by averaging the ΔF values from all reporter units (Hassan *et al*., 2021; **Figure 1D**). Pairs of units were included in the ΔF analysis if three criteria were met: (1) the PIC was fully activated, defined as the test unit being recruited at least 1s after the reporter unit (Bennett *et al*., 2001; Powers *et al*., 2008); (2) the test–reporter pair received common synaptic input, indicated by a rate–rate correlation of r ≥ 0.7 (Gorassini *et al*., 2004); and (3) the reporter unit exhibited a discharge range of at least 0.5 pulses per second (pps) while the test unit was active (Stephenson & Maluf, 2011).

A geometric measure, known as brace height (Beauchamp *et al*., 2023), was quantified to provide an additional measure of the contribution of PICs. Brace height was calculated as the maximal orthogonal deviation between the smoothed discharge rate profile of a motor unit and the predicted increase from recruitment to peak discharge rate. These values were normalized to the height of a right triangle (%rTri), where the hypotenuse represents recruitment to peak motor unit discharge (**Figure 1D**). The slopes of the initial acceleration and post-acceleration attenuation in discharge rate modulation of the motor unit discharge rate versus dorsiflexion torque were also quantified (Beauchamp *et al*., 2023).

### Statistical Analysis

To assess whether the distribution of menstrual cycle phases (early follicular, late follicular, mid luteal) was balanced across the three testing sessions, a chi-square test of independence was conducted. The analysis was based on each participant-phase-session combination to count the number of participants in each phase during each session. Linear mixed effects models were used to determine if the fixed effects of session or phase influenced MVC, with a random effect of participant.

### Menstrual cycle phase analysis

We used linear mixed-effects models to examine the effects of menstrual cycle phase and contraction intensity on each motor unit property, following the general model structure: Outcome ∼ Phase + Intensity + Session + Recruitment Threshold + (1 | Participant/Trial). The model was fit using the *lme4* package (Bates *et al*., 2015) in R (R Core Team, 2021) and assumptions were evaluated using the *Performance* package (Lüdecke *et al*., 2021). All variables met assumptions in their raw scale except acceleration and attenuation slopes, which showed non-normal residuals and required log transformation prior to fitting. Each model included the fixed effects of phase (early follicular, late follicular, mid luteal) and intensity (30, 50, and 70% MVC). Interactions between phase and intensity were explored by comparing models with and without the interaction term. The interaction was included in the final model when it significantly improved model fit according to log-likelihood ratio tests. Covariates for motor unit recruitment threshold and session (1, 2, 3) were included in the final models. Recruitment threshold was added to account for known associations between recruitment and motor unit discharge properties, and session number was included to control for potential learning effects across repeated visits. Both covariates significantly improved model fit based on log-likelihood ratio tests comparing models with and without each term. Random intercepts were specified for each participant (1 | participant) and for trials nested within participants (1 | Trial:Participant) to account for repeated measures. All categorical variables were coded as factors prior to model fitting. Each model was also weighted by the motor unit yield of each trial. Weighted models were selected over unweighted models based on both conceptual and statistical justification. Trials with higher motor unit yield provide more valid estimates of motor unit properties from the entire motor unit pool, such as ΔF, and the weighted models improved model fit, including lower AIC and BIC values and a significantly higher log-likelihood compared to unweighted models.

Estimated marginal means were calculated using the *emmeans* package in R (Lenth *et al*., 2023). Pairwise contrasts were performed to compare levels of phase and levels of intensity. When a phase by intensity interaction was present, simple effects were evaluated by estimating marginal means within each level of the interacting factor. Tukey adjustment was applied for all post hoc comparisons. Estimated marginal means and pairwise contrasts across phases were computed using the *emmeans* package. Cohen’s d effect sizes were obtained from the model-based contrasts using the *eff_size()* function. Effect sizes are reported in text as absolute values. To ensure that fixed effects of phase and session could be interpreted independently, we assessed multicollinearity using generalized variance inflation factors (GVIFs) from the *car* package. Estimated marginal means and their 95% confidence intervals were then visualized using *ggplot2* (Wickham *et al*., 2023).

### Multilevel modeling of hormone effects

To examine how circulating sex hormones related to motor unit discharge properties, we used multilevel linear mixed-effects models implemented using *lme4* (Bates *et al*., 2015) and *lmerTest* (Kuznetsova *et al*., 2017) in R (R Core Team, 2021). Because estradiol and progesterone concentrations are right-skewed, both hormones were log-transformed prior to analysis to reduce influence of high values and to better approximate model normality. Hormones were modeled using a within-between decomposition, separating each participant’s average hormone level from their menstrual cycle deviations.

For each hormone, the participant-level mean was calculated as the average value across the three phases and was used to represent between-participant variation. Within-participant hormone fluctuations were then calculated by centering each phase-specific value around that participant-level mean. This mean-centered score captures within-participant hormonal changes across the menstrual cycle and allows the model to separately estimate within- and between-participant hormonal effects.

Separate multilevel models were run for peak discharge rate, brace height, and ΔF using the general model structure: Outcome ∼ P4 mean + P4 within + E2 mean + E2 within + Intensity + Recruitment Threshold + (1 | Participant/Trial). All models included participant-level hormone means and within-participant hormone fluctuations for each hormone and contraction intensity as fixed effects, and recruitment threshold as a covariate. Repeated measures were modeled using a nested random intercept to account for trials nested within participants, and all models were weighted by motor unit yield. For peak discharge rate only, adding an interaction between hormone and intensity improved model fit based on a log-likelihood test.

Predicted values from the multilevel models were generated using the *ggeffects* (Lüdecke *et al*., 2025) package in R, which provides marginal predictions from mixed-effects models while holding other covariates at their observed or mean values. For each outcome, *ggpredict()* was used to obtain estimated marginal slopes for within-participant hormone fluctuations at each contraction intensity. To allow comparison of hormone effect sizes across outcomes with different units and scales (ΔF, peak discharge rate, and brace height), a subsequent analysis was run where each outcome was standardized (z-scored) prior to modeling. This transformation does not change the pattern of results but expresses effects in standard deviation units, facilitating direct cross variable comparison.

## RESULTS

### Participants

Across all sites, nine participants were excluded based on hormonal criteria, and one was excluded due to the absence of valid motor unit decomposition results, resulting in a final sample size of 40 females which were included in subsequent analyses (see Figure 2 for hormone concentrations). Of those included, not all participants completed ramp contractions at every target intensity: 40 completed ramps at 30% MVC, 31 at 50% MVC, and 30 at 70% MVC.

**Figure 2.**
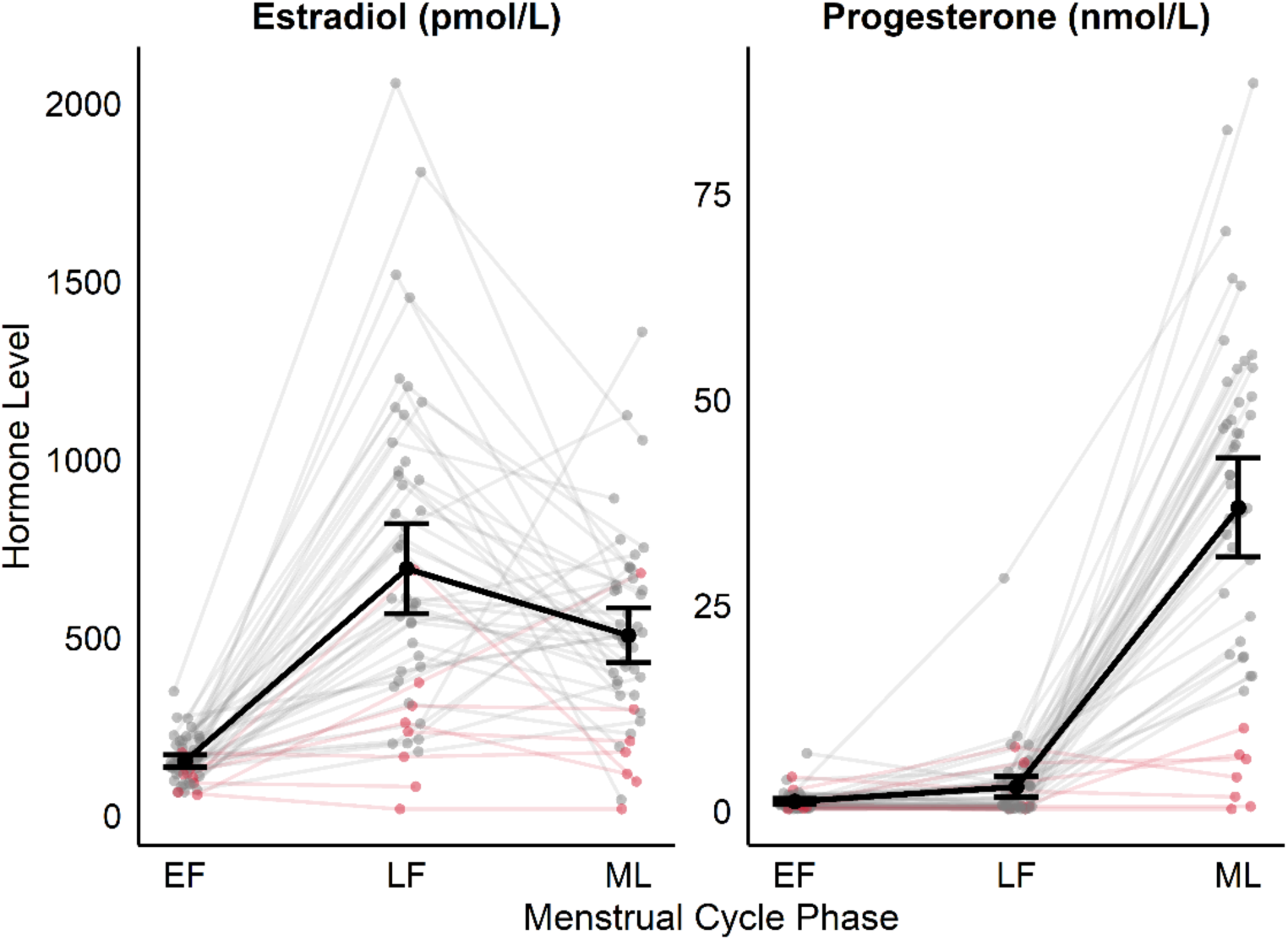
Individual dots and lines represent hormone levels for each participant at each phase of the cycle, pink participants were excluded due to no positive ovulation test during testing cycle or progesterone levels below 12 nmol/L in the ML phase.

### Cycle characteristics

The average cycle duration of participants’ tracked cycles was 27 ± 3 days. Testing sometimes occurred across multiple cycles due to randomized session order, so the average cycle length for participants during testing was 29 ± 2 days. On average, the early follicular visit was performed on day 3 ± 1, the late follicular visit on day 13 ± 3, and the mid luteal visit 7 ± 2 days after confirmed ovulation. Ovulation was confirmed in all the tracked cycles, occurring on day 14 ± 3. In the test cycle(s) used for laboratory visits, the positive LH surge test typically occurred 2 ± 2 days after the late follicular visit, consistent with typical timing of ovulation in eumenorrheic adults.

### Session randomization

Among the included participants, the onset of the menstrual cycle was sufficiently randomized across sessions. Fifteen participants began their cycle (early follicular phase) at session 1, thirteen at session 2, and twelve at session 3. A chi-square test revealed no significant association between cycle phase and session number (χ²(4) = 1.2, *p* = 0.878), indicating that menstrual phases were evenly distributed across sessions. This supports the effectiveness of randomized session scheduling with respect to cycle phase. MVC was similar at each session (χ²(1) = 0.08, *p* = 0.78) and was not influenced by menstrual cycle phase (χ²(2) = 0.21, *p* = 0.90; Table 1).

**Table 1.** Estimated marginal means (Est. Mean) and 95% confidence intervals (lower limit [LL] and upper limit [UL]) from the linear mixed effects of MVC across menstrual cycle phases, shown separately by site due to differences in measurement units (either in Newtons [N] or Newton-meters [Nm]).

| Site 1 |  |  |  | Site 2 |  |  |  | Site 3 |  |  |  |
| --- | --- | --- | --- | --- | --- | --- | --- | --- | --- | --- | --- |
| Phase | Est. Mean<br>(N) | LL | UL | Phase | Est. Mean<br>(Nm) | LL | UL | Phase | Est. Mean<br>(N) | LL | UL |
| EF | 16.7 | 14.6 | 18.8 | EF | 16.1 | 14.1 | 18.2 | EF | 14.3 | 12.5 | 16 |
| LF | 16.6 | 14.5 | 18.6 | LF | 16 | 13.9 | 18.1 | LF | 14.1 | 12.4 | 15.9 |
| ML | 16.7 | 14.6 | 18.7 | ML | 16.1 | 14 | 18.2 | ML | 14.3 | 12.5 | 16 |

**Peak discharge rate** was predicted by both menstrual cycle phase (χ²(2) = 31.56, *p* < 0.0001) and contraction intensity (χ²(2) = 1598.7, *p* < 0.0001). Post-hoc contrasts showed that peak discharge rate was lower in the early follicular phase than in both the late follicular (estimate = -0.59 ± 0.12, *p* < 0.0001, d =0.22) and mid luteal phases (estimate = -0.60 ± 0.12, *p* < 0.0001, d = 0.22), with no difference between late follicular and mid luteal phases (p = 0.10; Figure 3A). As expected, peak discharge rate increased progressively with contraction intensity, being lowest at 30% MVC and highest at 70% MVC. Estimated marginal means for each phase and intensity are shown in Table 2.

**Figure 3.**
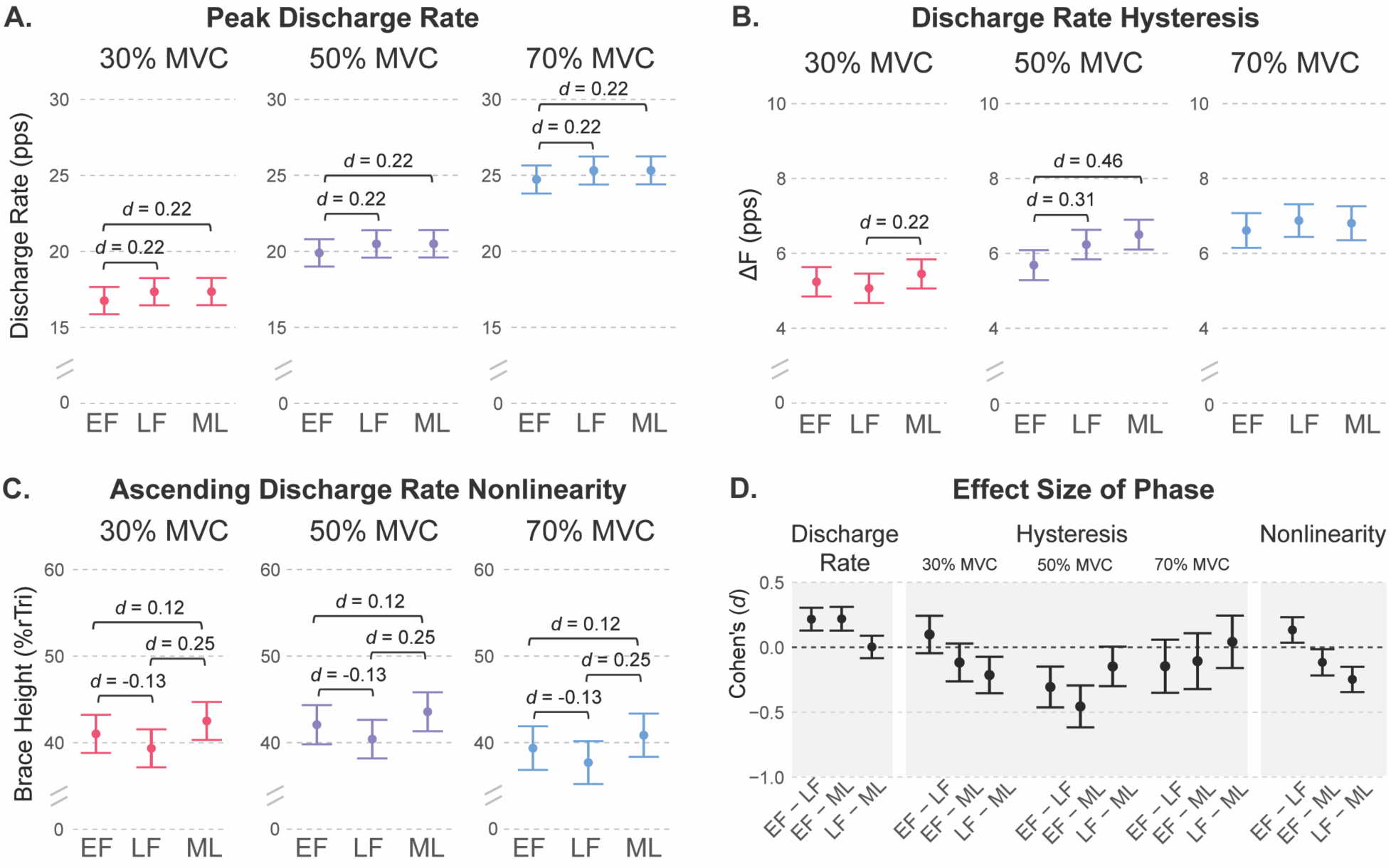
Estimated marginal means with 95% confidence intervals and effect sizes (d) for significant contrasts of (A) Peak discharge rate, (B) ΔF, and (C) Brace Height at each menstrual cycle phase. (D) Effect sizes for each measure, peak discharge rate and ascending discharge rate nonlinearity (Brace Height) across all intensities, and the interaction between phase and contraction intensity for discharge rate hysteresis (ΔF).

**Table 2.** Estimated marginal means (Est. Mean) and 95% confidence intervals (lower limit [LL] and upper limit [UL]) from the linear mixed effects models for all motor unit properties across phase and intensity.

| A. Peak Discharge Rate (pps) |  |  |  |  |  |  |  |  |  |  |  |
| --- | --- | --- | --- | --- | --- | --- | --- | --- | --- | --- | --- |
| 30% MVC |  |  |  | 50% MVC |  |  |  | 70% MVC |  |  |  |
| Phase | Est. Mean | 95% LL | 95% UL | Phase | Est. Mean | 95% LL | 95% UL | Phase | Est. Mean | 95% LL | 95% UL |
| EF | 16.8 | 15.9 | 17.7 | EF | 19.9 | 19 | 20.8 | EF | 24.7 | 23.8 | 25.7 |
| LF | 17.3 | 16.5 | 18.2 | LF | 20.5 | 19.6 | 21.4 | LF | 25.3 | 24.4 | 26.2 |
| ML | 17.4 | 16.5 | 18.3 | ML | 20.5 | 19.6 | 21.4 | ML | 25.3 | 24.4 | 26.2 |
| B. Discharge Rate Hysteresis ($\Delta F$; pps) | | | | | | | | | | | |
| 30% MVC |  |  |  | 50% MVC |  |  |  | 70% MVC |  |  |  |
| Phase | Est. Mean | 95% LL | 95% UL | Phase | Est. Mean | 95% LL | 95% UL | Phase | Est. Mean | 95% LL | 95% UL |
| EF | 5.24 | 4.85 | 5.63 | EF | 5.69 | 5.29 | 6.09 | EF | 6.61 | 6.15 | 7.08 |
| LF | 5.07 | 4.68 | 5.46 | LF | 6.23 | 5.84 | 6.63 | LF | 6.88 | 6.44 | 7.31 |
| ML | 5.45 | 5.06 | 5.84 | ML | 6.50 | 6.10 | 6.90 | ML | 6.81 | 6.35 | 7.26 |
| **C. Ascending Discharge Rate Nonlinearity (Brace Height; %rTri)** |  |  |  |  |  |  |  |  |  |  |  |
| **30% MVC** |  |  |  | **50% MVC** |  |  |  | **70% MVC** |  |  |  |
| **Phase** | **Est. Mean** | **95% LL** | **95% UL** | **Phase** | **Est. Mean** | **95% LL** | **95% UL** | **Phase** | **Est. Mean** | **95% LL** | **95% UL** |
| EF | 41.0 | 38.8 | 43.2 | EF | 42.1 | 39.8 | 44.3 | EF | 39.4 | 36.8 | 41.9 |
| LF | 39.3 | 37.1 | 41.5 | LF | 40.4 | 38.2 | 42.6 | LF | 37.7 | 35.2 | 40.2 |
| ML | 42.5 | 40.3 | 44.7 | ML | 43.6 | 41.3 | 45.8 | ML | 40.9 | 38.4 | 43.4 |
| **D. Acceleration Slope (pps/%MVC)** |  |  |  |  |  |  |  |  |  |  |  |
| **30% MVC** |  |  |  | **50% MVC** |  |  |  | **70% MVC** |  |  |  |
| **Phase** | **Est. Mean** | **95% LL** | **95% UL** | **Phase** | **Est. Mean** | **95% LL** | **95% UL** | **Phase** | **Est. Mean** | **95% LL** | **95% UL** |
| EF | 2.14 | 1.95 | 2.34 | EF | 1.27 | 1.16 | 1.39 | EF | 0.80 | 0.72 | 0.90 |
| LF | 2.20 | 2.03 | 2.43 | LF | 1.32 | 1.28 | 1.44 | LF | 0.83 | 0.75 | 0.92 |
| ML | 2.39 | 2.18 | 2.61 | ML | 1.42 | 1.30 | 1.56 | ML | 0.90 | 0.81 | 0.99 |
| **E. Attenuation Slope (pps/%MVC)** |  |  |  |  |  |  |  |  |  |  |  |
| **30% MVC** |  |  |  | **50% MVC** |  |  |  | **70% MVC** |  |  |  |
| **Phase** | **Est. Mean** | **95% LL** | **95% UL** | **Phase** | **Est. Mean** | **95% LL** | **95% UL** | **Phase** | **Est. Mean** | **95% LL** | **95% UL** |
| EF | 0.32 | 0.29 | 0.35 | EF | 0.18 | 0.16 | 0.20 | EF | 0.14 | 0.12 | 0.16 |
| LF | 0.35 | 0.31 | 0.38 | LF | 0.19 | 0.17 | 0.21 | LF | 0.15 | 0.14 | 0.17 |
| ML | 0.31 | 0.30 | 0.37 | ML | 0.18 | 0.16 | 0.20 | ML | 0.15 | 0.13 | 0.17 |

### Discharge rate hysteresis

There was a significant interaction between menstrual cycle phase and contraction intensity for ΔF (χ²(8) = 157.58, *p* < 0.0001). At 30 % MVC, ΔF was greater in the mid luteal compared with late follicular phase (estimate = -0.38 ± 0.12, *p* = 0.006, d = 0.21), with no difference between early follicular and either late follicular (*p* = 0.37, d = 0.10) or mid luteal (*p* = 0.23, d = 0.12) phases. At 50% MVC, ΔF was greater in the mid luteal than early follicular (estimate = -0.81 ± 0.14, *p* < 0.0001, d = 0.46) and greater in the late follicular than early follicular phase (estimate = -0.55 ± 0.14, *p* = 0.0002, d = 0.31), but similar at late follicular and mid luteal (estimate = -0.27 ± 0.13, *p* = 0.12, d = 0.15). At 70% MVC, ΔF did not differ between phases (*p* > 0.32 for all comparisons). Estimated marginal means for each phase and intensity are presented in Table 2, and the interaction is illustrated in Figure 3.

**Ascending discharge rate nonlinearity** was quantified using the brace height metric as well as acceleration and attenuation slopes. Brace height was influenced by both menstrual cycle phase (χ²(2) = 26.88, *p* < 0.0001) and contraction intensity (χ²(2) = 12.85, *p* = 0.002). Estimated marginal means showed a consistent pattern across intensities. From early follicular to late follicular, brace height decreased (estimate = -1.67 ± 0.62, *p* = 0.019; d = 0.13) and then increased from late follicular to mid luteal (estimate = 3.16, SE = 0.61, *p* < 0.001; d = 0.25). This was the largest difference, although the effect size was still small to moderate. The mid luteal phase was also higher than early follicular (estimate = 1.49, SE = 0.64, *p* = 0.050; d = 0.12).

Acceleration and attenuation slopes were analyzed on the log scale due to improved model fit and residual normality following log transformation. Both phase (χ²(2) = 15.169, *p* < 0.001) and contraction intensity (χ²(2) = 442.47, *p* < 0.0001) were significant predictors of acceleration slope. Across intensities, acceleration slope was greater in the mid luteal compared with both early follicular (ratio = 0.89 ± 0.02, *p* < 0.001, d = 0.19) and late follicular phases (ratio = 0.92 ± 0.03, *p* = 0.005, d = 0.14). Estimated geometric means for each phase and intensity are presented in Table 2D, and the consistent phase effects are illustrated in Figure 3.

Only contraction intensity was a significant predictor of attenuation slope (χ²(2) = 288.49, *p* < 0.0001), with progressively lower values at higher intensities. Menstrual cycle phase showed a modest effect (χ²(2) = 5.85, *p* = 0.054), driven by a slightly lower slope in the early follicular compared with late follicular phase (ratio = -0.92 ± 0.03, *p* = 0.042, d = 0.11). No differences were observed between the early follicular and mid luteal (*p* = 0.46) or late follicular and mid luteal phases (*p* = 0.45). Estimated means for each phase and intensity are presented in Table 2.

### Effects of session order

As mentioned previously, session number was included in the models because it improved model fit and accounted for potential learning effects across sessions. Adding session significantly improved model fit for peak discharge rate (χ²(2) = 123.15, *p* < 0.0001), ΔF (χ²(2) = 30.70, *p* < 0.0001), and brace height (χ²(2) = 7.21, *p* = 0.027), but not for acceleration slope (χ²(2) = 5.80, *p* = 0.055) or attenuation slope (χ²(2) = 0.39, *p* = 0.823). Importantly, for the motor unit discharge properties where both session and menstrual cycle phase were significant fixed effects (peak discharge rate, ΔF, and brace height), collinearity diagnostics confirmed that these predictors could be interpreted independently. Scaled generalized variance inflation factors (GVIFs) were low (all < 2), including values of 1.03 for phase and 1.03 for session in the peak discharge rate model, 1.55 for phase and 1.04 for session in the ΔF model, and 1.03 for phase and 1.03 for session in the brace height model. These results indicate that changes across menstrual cycle phases reflect true cycle effects rather than artifacts of session order or learning.

### Relationships between hormone concentrations and motor unit discharge

When we decomposed each hormone into within-person (cycle-related) and between-person (average) components, within-person estradiol showed significant associations with peak discharge rate and brace height, whereas within-person progesterone was associated with peak discharge rate, ΔF and brace height (Figure 4A-C).

**Figure 4.**
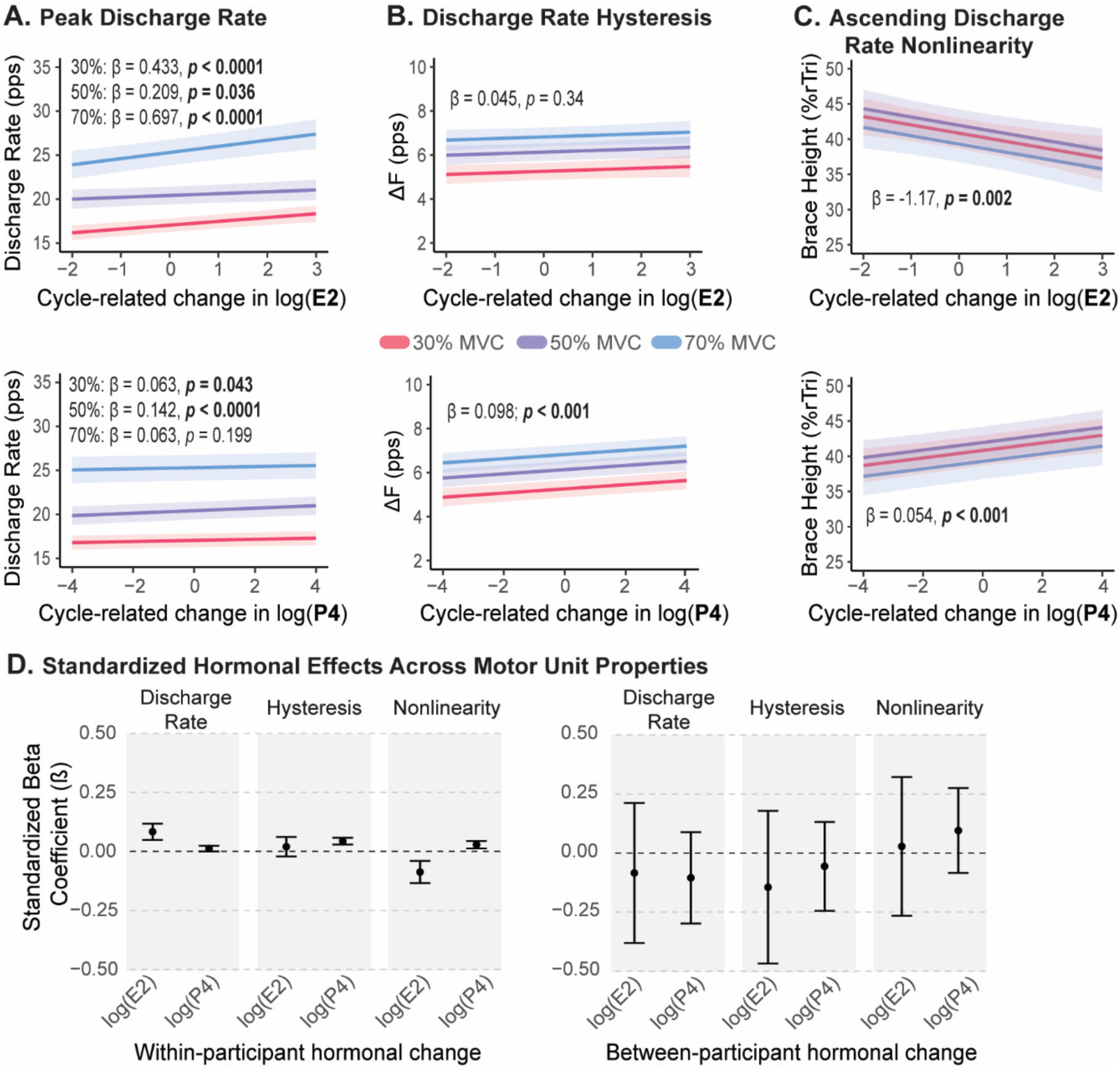
Within-person associations between cycle-related hormone fluctuations and motor-unit outcomes for (A) peak discharge rate, (B) discharge rate hysteresis, and (C) discharge rate nonlinearity. Each panel shows model-predicted values at each contraction intensity from linear mixed-effects models in which within-person hormone values (mean-centered log-transformed estradiol or progesterone) were used as predictors. “Cycle-related change” reflects the deviation of each observation from an individual’s mean log hormone level across sessions, with estradiol shown in the first row and progesterone shown in second. The bottom panel (D) shows standardized beta coefficients for the within- and between- participant hormone main effects.

The model for peak discharge rate showed significant main effects of both hormones within-individuals. Higher estradiol within an individual showed a positive association with peak discharge rate (β = 0.433 ± 0.091, *p* < 0.0001), and higher progesterone within an individual was associated with a modest increase (β = 0.063 ± 0.031, *p* = 0.043). Neither between-person estradiol (β = -0.47 ± 0.78, p = 0.55) or between person progesterone (β = -0.58 ± 0.51, *p* = 0.26) predicted peak discharge rate, indicating that females with higher overall hormone levels did not systematically differ from those with lower levels. Although the model with the best fit included hormone and intensity interactions, none of these interaction terms reached statistical significance, suggesting that the magnitude of the progesterone- or estradiol-related change in peak discharge rate did not differ across contraction intensities. Model predicted values at each intensity are show in in Figure 4A.

Increases in progesterone within an individual were associated with significantly higher ΔF (β = 0.098 ± 0.016, *p* < 0.0001), indicating that discharge rate hysteresis increased when progesterone rose above a participant’s own mean across the cycle (Figure 4B). The between-person progesterone effect was not significant (β = -0.14 ± 0.22, *p* = 0.51), showing no evidence that females with higher overall progesterone levels differed in ΔF from females with lower average levels. Estradiol was not a significant predictor of ΔF in terms of within-person estradiol fluctuations (β = 0.045 ± 0.047, *p* = 0.34) or between-person estradiol differences (β = -0.34 ± 0.37, *p* = 0.36). Together, these findings suggest that ΔF is specifically sensitive to within-person changes in progesterone concentrations.

Both hormones showed within-person associations with ascending discharge rate nonlinearity (i.e., brace height; Figure 4C). Higher estradiol within an individual was associated with slightly smaller brace height values (β = -1.17 ± 0.37, *p* = 0.002), whereas higher progesterone concentrations within an individual were associated with slightly higher brace height values (β = 0.54 ± 0.13, *p* < 0.001). In contrast, brace height was not predicted by between-person estradiol (β = - 0.97± 2.48, *p* = 0.70) or between-person progesterone (β = 1.6 ± 1.51, *p* = 0.29).

To determine whether hormonal fluctuations exert stronger effects on some motor unit properties than others, the outcomes were Z-scored to allow direct comparison across variables (Figure 4D). In this framework, coefficients represent standardized effect sizes in units of the outcome’s standard deviation. All estimates fell below 0.1 standard deviations, a range typically interpreted as a small effect. For peak discharge rate, higher estradiol was associated with a slightly larger increase (estimate = 0.08 ± 0.018, *p* < 0.001), and progesterone was associated with a small increase (estimate = 0.01 ± 0.006, *p* = 0.043). Estradiol showed no significant effect on ΔF (estimate = 0.02 ± 0.021, *p* <0.34), but progesterone was associated with greater discharge rate hysteresis (estimate = 0.04 ± 0.007, *p* < 0.0001). For brace height, higher estradiol was associated with decreases (estimate = -0.09 ± 0.024, *p* < 0.001), whereas higher progesterone was associated with increases in brace height (estimate = 0.03 ± 0.008, *p* < 0.001). Hormone-by-intensity interactions were not significant for any outcome. Again, the average level of both estradiol and progesterone between participants was not significant (shown in figure 4D). Overall, these findings indicate that within-participant hormonal fluctuations exert detectable but relatively small effects on motor unit behavior, and that these effects are similar in magnitude across different motor unit properties.

## DISCUSSION

The findings of this study show that motor unit discharge properties vary across the menstrual cycle, and these variations are associated with within-participant changes in estradiol and progesterone concentrations. Although detected changes were small, estradiol had a stronger influence, with higher levels associated with small increases in peak discharge rates and more linear ascending discharge rate patterns. Progesterone showed slightly weaker relationships, with higher concentrations associated with small increases in peak discharge rate, discharge rate hysteresis, and ascending discharge rate nonlinearity. Although these differences were statistically significant and therefore detectable, effect sizes were small. This emphasizes the importance of large sample size research with rigorous hormonal inclusion criteria. The absence of changes in MVC further suggests that cycle-related hormonal shifts likely do not affect gross measures of strength, aligning with meta-analyses of performance outcomes (de Jonge, 2003; McNulty *et al*., 2020).

### Menstrual cycle characteristics

Of the 50 females who completed the full protocol, only 41 had hormonal profiles consistent with the targeted menstrual cycle phase classifications and therefore were included in final analyses. Achieving a sample of this size with verified hormone concentrations required coordinated, multi-site recruitment and rigorous participant tracking, reflecting one of the major methodological challenges of investigation into the menstrual cycle. Notably, approximately one in five participants exhibited hormone profiles that did not match the expected cycle phase based on calendar tracking alone. This proportion aligns with epidemiological evidence showing that up to a quarter (14-25%) of cycles in healthy, regularly menstruating adults are anovulatory or hormonally atypical (Whitaker & Critchley, 2016). These patterns are physiologically common, underscoring that even among individuals with regular self-reported cycles and prospective tracking, menstrual cycle phase cannot be assumed without hormone verification. This has direct implications for the interpretation of prior literature. Many of the earlier studies comparing menstrual cycle phases included small samples (often 8-12 participants) and relied on self-report or calendar counting without serum hormone verification. Under these conditions, it is highly likely that participants classified as being in *different phases*, may not have had the hormonal profile assumed for that phase. If phase effects were quantified, but hormone measures were not taken, it limits the ability to interpret direct hormonal effects. Our approach, which incorporated prior tracking, luteinizing hormone surge testing, and serum hormone measurement, ensured accurate phase classification and allowed cycle-related effects to be interpreted in relation to the underlying hormonal fluctuations. Furthermore, our subsequent analysis of the effects of estradiol and progesterone on motor unit properties helps the interpretation of phase differences. This does not imply that individuals with atypical or anovulatory cycles should never be studied, but it highlights that when the research question specifically concerns phase-based differences, hormonal verification is essential to ensure that phase designations reflect true physiological state. Future research incorporating both hormonally verified phases and characterization of atypical cycles are needed to advance mechanistic understanding of female neuromuscular physiology.

### Previous menstrual cycle studies

Other studies have documented changes in motor unit discharge across the menstrual cycle, but these have been limited to sustained, low intensity isometric contractions where only initial or average discharge rates were quantified. In the first study to examine menstrual cycle effects on motor unit behavior, Tenan et al. (2013) reported increases in initial discharge rate of single motor units recorded using fine-wire EMG in the vastus medialis across the menstrual cycle. Members of the MUSH collaboration have more recently shown menstrual cycle-related differences in average discharge rate during sustained isometric contractions to target forces of 10 and 25% MVC using intramuscular EMG in the vastus lateralis (Piasecki *et al*., 2023). However, these prior studies did not quantify properties that reflect the contribution of PICs on motor unit discharge patterns or explore intensities above 30% MVC. Previous work from our group and others have now established clear sex-related differences in estimates of PIC contributions to motor unit discharge, with females showing greater values than males (Jenz *et al*., 2023; Lecce *et al*., 2024; Yacyshyn *et al*., 2025). Several studies have also reported higher motor unit discharge rates during submaximal isometric contractions in females compared with males (Inglis & Gabriel, 2020; Guo *et al*., 2022; Taylor *et al*., 2022; Jenz *et al*., 2023). In most of these studies, the question of whether sex hormone concentrations affect motoneuron function has repeatedly been posed.

### Within-person hormonal changes predict changes in motor unit discharge

Changes in hormone levels within-participants across their menstrual cycle showed that progesterone and estradiol had divergent effects on the motor unit properties quantified. Higher levels of progesterone within individuals were associated with more discharge rate hysteresis, nonlinearity of the ascending phase, and higher peak discharge rates, whereas increases in estradiol was associated with higher peak discharge rates and more linear ascending discharge rates. Neither hormone demonstrated significant between-person effects, which suggests that absolute hormone levels were less informative than individual fluctuations within each participant’s typical physiological range. This is likely because the females included in this study had similar average hormone levels.

Future studies comparing the effects of these hormones in females with attenuated values due to hormonal contraceptives or menopause will provide more insight. Z-scored beta coefficients allowed for the analysis of which motor unit property was most greatly affected by within-cycle hormone changes. While most beta coefficients were lower than 0.1, indicating a relatively small effect compared to the standard variability, estradiol predicted peak discharge rate (β = 0.083) and ascending discharge rate nonlinearity (β = -0.87) stronger than any other significant effect.

These hormonal effects align with the differences detected across menstrual cycle phases, reaffirming the differences observed at each phase are hormone driven. Peak discharge rate, with a positive relationship with estradiol, increased from early follicular to late follicular, and remained elevated in the mid luteal phase, following the pattern of elevated estradiol in those later phases. Similarly, the negative relationship between estradiol and brace height concurs with the decrease in brace height in the late follicular phase. Although the positive relationship between progesterone and ascending discharge rate nonlinearity was weaker, it is still reflected in the elevation of brace height measures in the mid luteal phase, consistent with the rise in progesterone. Despite this, the change in discharge rate hysteresis is challenging to interpret due to differences across phases at different intensities. However, the weak but positive relationship between progesterone and ΔF suggests an effect on the neuromodulatory drive to motoneurons by progesterone, supported by differences in the mid luteal phase at 30 and 50% MVC.

### Linking Sex Hormones to Persistent Inward Currents

The effects of sex hormones on motor unit discharge patterns noted above likely relate to shifts in neuromodulatory and inhibitory inputs to motoneurons. PICs are facilitated by serotonin and norepinephrine, originating in the brainstem nuclei (Heckman & Enoka, 2012), and both estradiol and progesterone have been shown to influence these systems in reduced preparations. However, the direct effects of these hormones on spinal circuits and, more specifically, spinal motoneurons in reduced preparations are not well-studied.

The effects of progesterone have mostly been documented as net inhibitory in several brain circuits (Smith *et al*., 1989; Hsu & Smith, 2003). Evidence from primate studies, however, showed that applying progesterone in an estradiol-primed state, mimicking the mid luteal phase, increases extracellular serotonin concentrations in the hypothalamus (Centeno *et al*., 2007). Indeed, serotonergic neurons in the dorsal and caudal raphe express the progesterone receptor (Bethea, 1993), which suggests that it is possible for a progesterone surge in the mid luteal phase to enhance serotonin release from the brainstem, which would result in facilitation of discharge rate hysteresis (i.e., ΔF) and ascending discharge rate nonlinearity (i.e., brace height).

Estradiol also has well-documented effects on serotonergic signaling. Estrogen receptor beta is expressed in serotonin-producing neurons (Gundlah *et al*., 2000), and depletion of estradiol reduces the number of these neurons (Bethea *et al*., 2011). In rats, restoring estradiol increases expression of the 5HT2A receptor (Sumner & Fink, 1993), which is also present on motoneuron dendrites, and 5HT2A receptor density naturally increases during the endogenous estradiol surge (Sumner & Fink, 1997). These hormonal influences on serotonergic signaling align with evidence of enhanced plateau potentials during estrus in anesthetized cats (Kirkwood *et al*., 2002; Ford & Kirkwood, 2006), suggesting increased PIC activation when hormones are elevated.

Increases in estradiol have also shown net excitation in neuronal signaling, either through direct influence on glutamatergic neurons (Smith *et al*., 1987, 1988) or through decreases in GABA-A mediated inhibition (Mukherjee *et al*., 2017). Again, these prior investigations have only shown net excitation in the brain. Therefore, the effect on spinal cord circuitry might be different, potentially having influences through the net excitation of inhibitory pathways, for example, the corticospinal neurons targeting presynaptic interneurons that inhibit spinal motoneurons or corticoreticular neurons. Increases in peak discharge rate with estradiol may reflect the overall net excitatory effect, while simultaneously the decrease in discharge rate nonlinearity may be due to altered inhibitory patterns, which may be influenced by estradiol. Together, this prior literature provides a plausible physiological mechanism for the effects of estradiol and progesterone on motor unit discharge observed in this study, but further work is required to draw more definitive conclusions.

## CONCLUSION

Differences in motor unit discharge behavior between menstrual cycle phases is related to within-participant fluctuations in estradiol and progesterone concentrations. Elevated estradiol and progesterone levels in the late follicular and mid luteal phases, respectively, align with higher peak discharge rates; increases in progesterone during the mid luteal phase align with greater discharge rate hysteresis; and higher estradiol levels in the late follicular phase predict more linear ascending discharge patterns, whereas higher progesterone during the mid luteal phase aligns with greater nonlinearity. Although effect sizes for phase-based motor unit metrics are small, within-participant hormonal fluctuations consistently reflect these modulations in motor unit discharge, supporting hormonal mechanisms underpinning these phase effects. Together, these findings suggest that endogenous fluctuations of sex hormones influence biophysical motoneuron properties and/or their inputs that govern motor unit discharge. A key strength of the MUSH collaboration is that its large sample size and methodological rigor enabled detection of these subtle effects. To determine whether the menstrual cycle meaningfully influences motor behavior and its underlying physiology, such experimental designs are essential.

## FUNDING

S.T.J. - National Institutes of Health grant [FAIN: F31NS130767]

J.P. & P.A - Royal Society Research Grant [RGS/R1/231080]

G.E.P.P. – Natural Sciences and Engineering Research Council of Canada – Discovery Grant [2023-05862]; Banting Discovery Foundation – Ontario Brain Institute Discovery Award; & Azrieli Foundation – Collaboration on Motor Planning, Execution and Resilience Network Grant

---

**Table A1.**
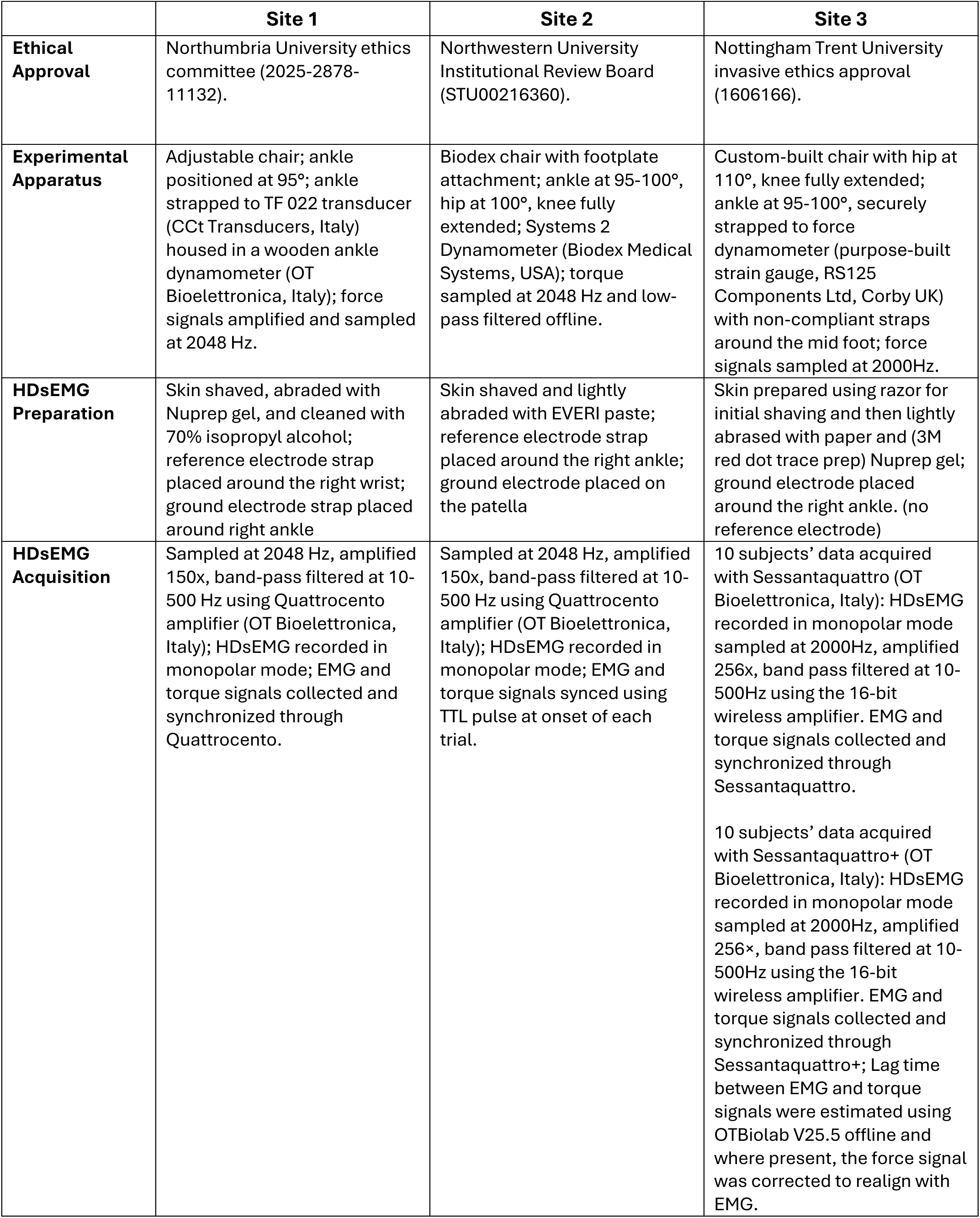
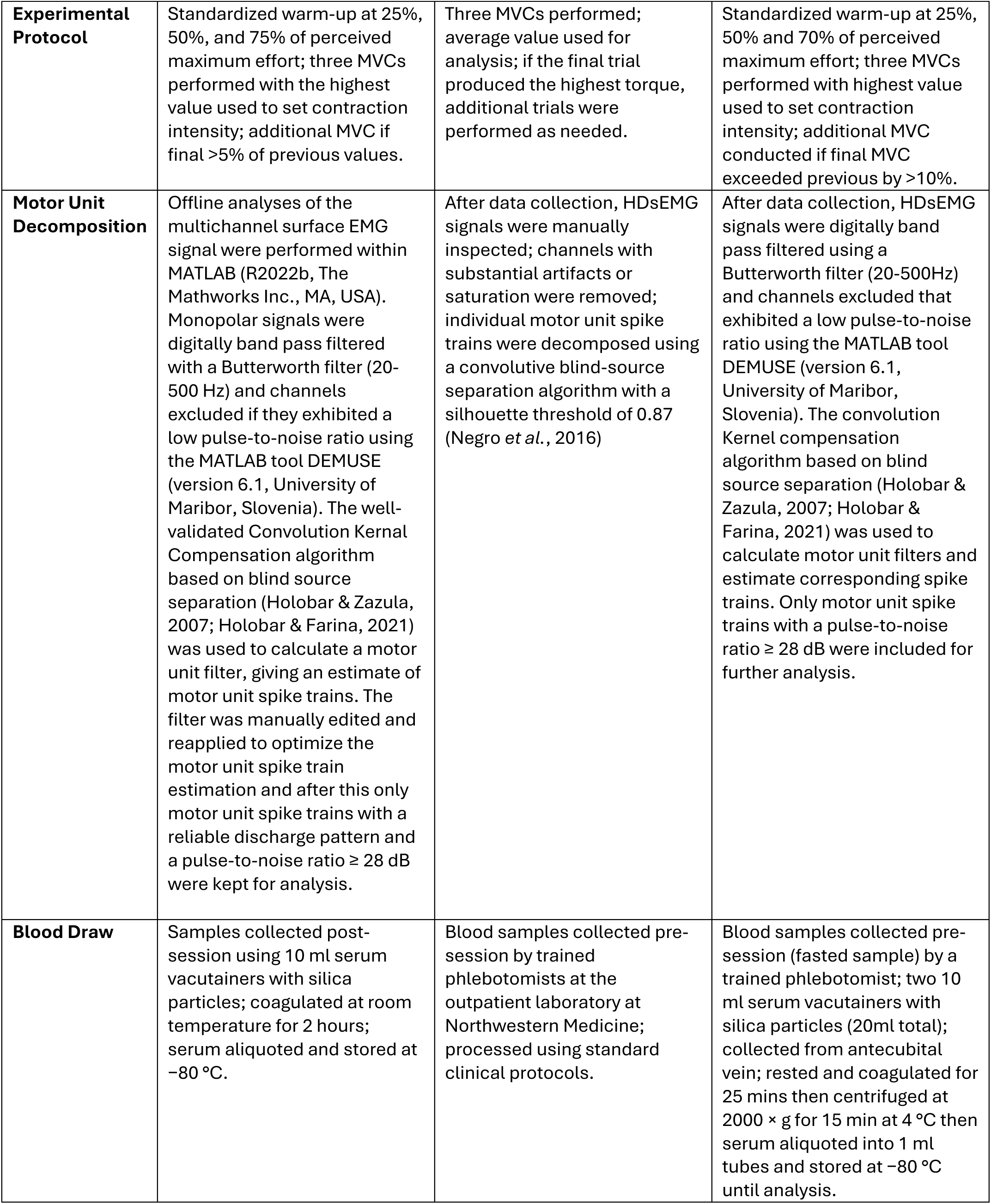
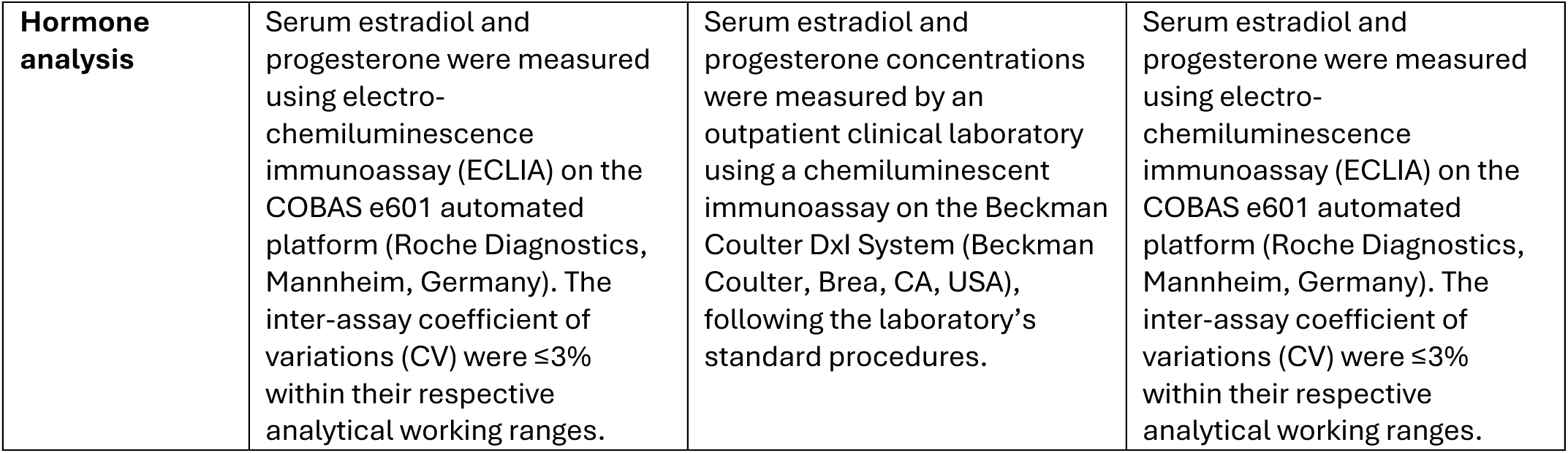
Experimental setup, HDsEMG procedures, motor unit decomposition methods, and hormone analysis across each testing site.

## Notes

### Competing Interest Statement

The authors have declared no competing interest.

